# Efficient genome editing in primary cells and *in vivo* using viral-derived “Nanoblades” loaded with Cas9/sgRNA ribonucleoproteins

**DOI:** 10.1101/202010

**Authors:** Philippe E. Mangeot, Valérie Risson, Floriane Fusil, Aline Marnef, Emilie Laurent, Juliana Blin, Virginie Mournetas, Emmanuelle Massouridès, Thibault J. M. Sohier, Antoine Corbin, Fabien Aube, Christian Pinset, Laurent Schaeffer, Gaëlle Legube, François-Loïc Cosset, Els Verhoeyen, Théophile Ohlmann, Emiliano P. Ricci

## Abstract

Programmable nucleases have enabled rapid and accessible genome engineering in eukaryotic cells and living organisms. However, their delivery into target cells can be technically challenging when working with primary cells or *in vivo*. Using engineered murine leukemia virus-like particles loaded with Cas9/sgRNA ribonucleoproteins (“Nanoblades”), we were able to induce efficient genome-editing in cell lines and primary cells including human induced pluripotent stem cells, human hematopoietic stem cells and mouse bone-marrow cells. Transgene-free Nanoblades were also capable of *in vivo* genome-editing in mouse embryos and in the liver of injected mice. Nanoblades can be complexed with donor DNA for “all-in-one” homology-directed repair or programmed with modified Cas9 variants to mediate transcriptional up-regulation of target genes. Nanoblades preparation process is simple, relatively inexpensive and can be easily implemented in any laboratory equipped for cellular biology.

## Introduction

Targeted genome editing tools such as Meganucleases (MGN), Zinc-finger nucleases (ZFN), Transcription Activator-Like Effector Nucleases (TALENs) and more recently the Clustered Regularly Interspaced Short Palindromic Repeats (CRISPR) have revolutionized most biomedical research fields. Such tools allow to precisely edit the genome of eukaryotic cells by inducing double-stranded DNA (dsDNA) breaks at specific loci. Relying on the cell endogenous repair pathways, dsDNA breaks can then be repaired by Non-Homologous End-Joining (NHEJ) or Homology-Directed Repair (HDR) allowing the removal or insertion of new genetic information at a desired locus.

Among the above mentioned tools, CRISPR/Cas9 is currently the most simple and versatile method for genome engineering. Indeed, in the two-component system, the bacterial derived nuclease Cas9 (for CRISPR-associated protein 9) associates with a single-guide RNA (sgRNA) to target a complementary DNA sequence and induce a dsDNA break^1^. Therefore, by the simple modification of the sgRNA sequence, users can specify the genomic locus to be targeted. Consistent with the great promises of CRISPR/Cas9 for genome engineering and gene therapy, considerable efforts have been made in developing efficient tools to deliver the Cas9 and the sgRNA into target cells *ex vivo* either by transfection of plasmids coding for the nucleases, transduction with viral-derived vectors coding for the nucleases or by direct injection or electroporation of Cas9/sgRNA complexes into cells.

Here, we have designed “Nanoblades”, a protein-delivery vector based on Friend Murine Leukemia Virus (MLV) that allows the transfer of Cas9/sgRNA ribonucleoproteins (RNPs) to cell lines, primary cells *in vitro* and *in vivo*. Nanoblades deliver the ribonucleoprotein cargo in a transient and rapid manner without delivering a transgene and can mediate knock-in in cell lines when complexed with a repair template. Nanoblades can also be programmed with modified Cas9 proteins to mediate transient transcriptional activation of targeted genes.

## Results

### Rapid and efficient delivery of Cas9/sgRNA RNPs through Murine Leukemia viruslike particles

Assembly of retroviral particles relies on the viral structural Gag polyprotein, which multimerizes at the cell membrane and is sufficient, when expressed in cultured cells, to induce release of virus-like particles (VLPs) into the cell supernatant ^2^. When Gag is coexpressed together with a fusogenic viral envelope, pseudotyped VLPs are produced that lack a viral genome but still retain their capacity to fuse with target cells and deliver the Gag protein into their cytoplasm. As previously investigated ^3^, we took advantage of the structural role of Gag and designed an expression vector coding for the MLV Gag polyprotein fused, at its C-terminal end, to a flag-tagged version of *Streptococcus pyogenes* Cas9 protein (Gag::Cas9, Figure 1a). The two fused proteins are separated by a proteolytic site which can be cleaved by the MLV protease to release the Flag-tagged Cas9 (Figure 1a). By cotransfecting HEK-293T cells with plasmids coding for Gag::Cas9, Gag-Pro-Pol, a single-guide RNA (sgRNA), and viral envelopes, fusogenic VLPs are produced and released in the culture medium (herein described as “Nanoblades”). Biochemical and imaging analysis of purified particles (Supplementary Figure 1a, 1b, 1c and 1d) indicates that Nanoblades (150nm) are slightly larger than wild-type MLV (Supplementary Figure 1b) but sediment at a density of 1.17 g/ml (Supplementary Figure 1c) as described for MLV VLPs ^4^. As detected by Western blot, Northern blot, mass-spectrometry and deep-sequencing, Nanoblades contain the Cas9 protein and sgRNA (Supplementary Figure 1 and 2 and Supplementary Table 1). Interestingly, the packaging of sgRNA depends on the presence of the Gag::Cas9 fusion protein since Nanoblades produced from cells that only express the Gag protein fail to incorporate detectable amounts of sgRNA (Supplementary Figure 1d).

**Figure 1.**
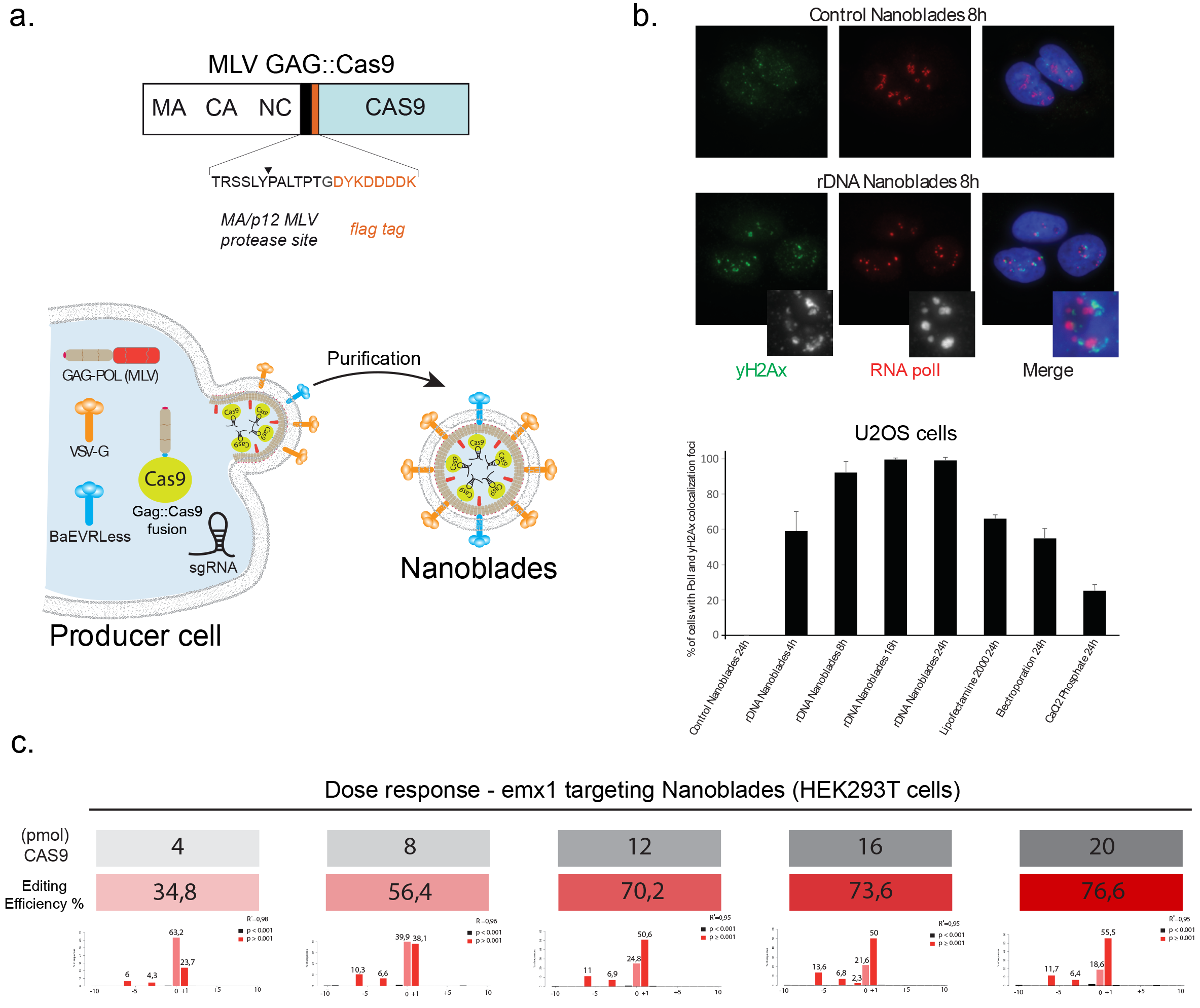
Nanoblade mediated genome editing. **a**. Scheme describing the MLV Gag::Cas9 fusion and the Nanoblade production protocol based on the transfection of HEK-293T cells by plasmids coding for Gag-Pol, Gag::Cas9, VSV-G, BaEVRLess and the sgRNA. Right panel, electron microscopy analysis of purified Nanoblades. **b**. Top panel, immunofluorescence analysis of γ-H2AX (green), RNA poll (red) in U2OS cells 8 hours after being transduced with control Nanoblades or with Nanoblades targeting ribosomal DNA genes. Bottom panel, quantification of γ-H2AX and RNA poll colocalization foci in U2OS cells at different times after Nanoblades transduction or after classical DNA transfection methods. **c**. Dose response of Nanoblades. HEK-293T cells were transduced with increasing amounts of Nanoblades targeting human *EMX1*. The exact amount of Cas9 used for transduction was measured by dot blot (in grey). Genome editing was assessed by Sanger sequencing and Tide analysis (in red).

To assess for Cas9/sgRNA RNP delivery efficiency in target cells and induction of genomic dsDNA breaks, we designed Nanoblades with a sgRNA targeting the 45S rDNA loci. Human rDNA genes are present in hundreds of tandem repeats across 5 autosomes, locate in the nucleolus and are transcribed exclusively by RNA polymerase (Pol) I ^5^. Using immunofluorescence microscopy, it is therefore possible to follow the occurrence of dsDNA breaks at rDNA loci with single-cell resolution by monitoring the nucleolus using the nucleolar marker RNA Pol I and the well-established dsDNA break-marker, histone variant γ-H2AX, that localizes at the nucleolar periphery after dsDNA break induction within rDNA ^6^. U2OS (osteosarcoma cell line) cells transduced for 24 hours with Nanoblades programmed with a sgRNA targeting rDNA display the typical γ-H2AX distribution at the nucleolar periphery with RNA Pol I, indicative of rDNA breaks, whilst cells transduced with Nanoblades with control sgRNAs do not (Figure 1b, top panel). Interestingly, this distribution of γ-H2AX at the nucleolar periphery can be observed as early as 4 hours after transduction in 60% of cells with a maximum effect observed at 16 hours after transduction, where almost 100% of observed cells display this γ-H2AX distribution (Figure 1b, bottom panel and quantification below). In comparison, only 60% of cells (at best) transfected with a plasmid coding for Cas9 and the sgRNA display the perinucleolar γ-H2AX/RNA Pol I localization 24 hours after transfection. Similar results were obtained in human primary fibroblasts with more than 85% cells displaying this distribution after 16 hours (Supplementary Figure 1e). These results suggest that Nanoblade-mediated delivery of the Cas9/sgRNA RNP is both efficient and rapid in cell lines and primary human cells.

To further confirm these results, we designed and dosed Nanoblades programmed with a sgRNA widely used in the literature that targets the human *EMX1* gene to induce a single cleavage. Indeed, loading of Cas9 into Nanoblades can be monitored from a small aliquot of Nanoblades by performing dot blot or elisa assays using anti Cas9 antibodies (Data not shown). This approach allows an indirect measure of viral titers to normalize the amount of Cas9 protein added to target cells. We then transduced HEK-293T cells with increasing amounts of Nanoblades and measured gene editing from the bulk population 48 hours after transduction (Figure 1c). Under these conditions, we observed a dose-dependent effect of Nanoblades ranging from 35% of *EMX1* editing to 77% of editing at the highest dose of Cas9 (Figure 1c).

### Nanoblade-mediated genome editing in human and mouse primary cells

Genome editing in primary cells and patient-derived pluripotent cells represents a major interest both for basic science and therapeutical applications. However, primary cells are often refractory to DNA transfection and other gene delivery methods. Because Nanoblades are capable of efficient delivery of functional Cas9/sgRNA RNPs into primary fibroblasts, we tested whether they were effective in other primary cell for genome editing. To this aim, Nanoblades targeting *EMX1* were used to transduce human induced Pluripotent Stem Cells (hiPSCs). Genome editing at the *EMX1* locus was assessed in the bulk cellular population 48 hours after transduction by deep-sequencing of the *EMX1* locus (Figure 2a, left panel). As observed, Nanoblades are capable of mediating 67% genome editing at the *EMX1* locus in hiPSCs. Similar results were obtained using iPS cells derived from a patient with Duchenne muscular dystrophy bearing a mutation in exon 7 transduced with Nanoblades programmed with a single sgRNA targeting the mutated loci. In this case, we obtained close to 40% of gene editing (Figure 2a, right panel). Notably, hiPSCs treated with *EMX1* Nanoblades maintained constant levels of pluripotency markers compared to control cells (Figure 2a, middle panel) thus indicating that their multipotent status does not appear to be affected.

**Figure 2.**
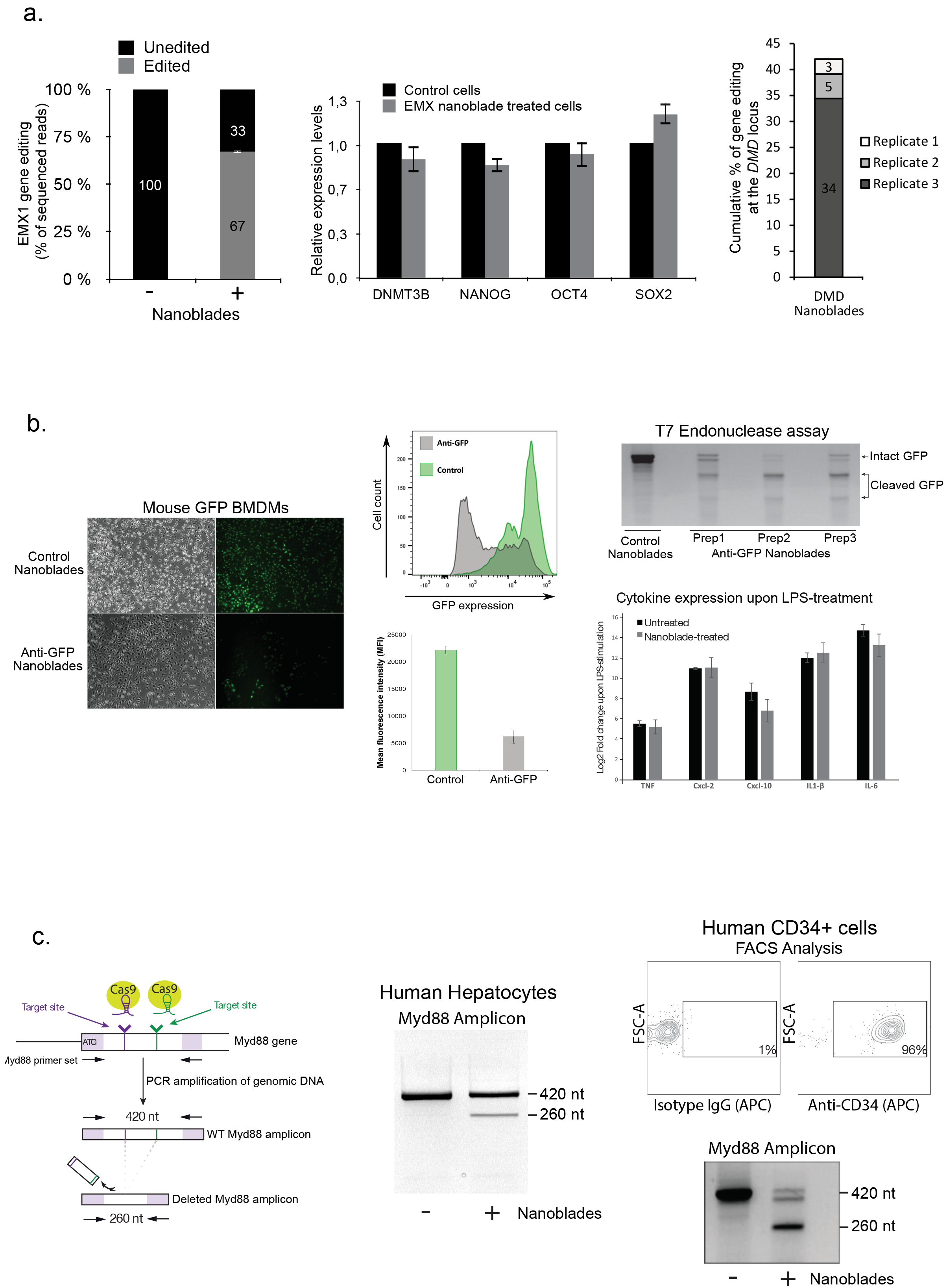
Genome editing in primary cells transduced with Nanoblades. **a**. Left panel, editing efficiency at the *EMX1* locus (measured by high-throughput sequencing on the Illumina Miseq platform) of human induced pluripotent stem cells (hiPSCs) transduced with Nanoblades targeting human *EMX1*. Middle panel, expression of pluripotency markers measured by qPCR in control cells and cells transduced with Nanoblades targeting *EMX1*. Right panel, editing efficiency at the *DMD* locus in hiPSCs derived from a patient with Duchenne muscular dystrophy transduced with Nanoblades targeting the mutated locus. **b**. Left and middle panels, fluorescence microscopy and FACS analysis of GFP expressing BMDMs transduced at the bone marrow stage (day 0 after bone marrow collection) with control Nanoblades or Nanoblades targeting the *GFP* coding sequence. Right panel, cytokine expression levels (measured by qPCR) in untreated or Nanoblade-treated cells upon LPS stimulation. **c**. Left panel, excision of a 160bp DNA fragment of *MYD88* using Nanoblades. Middle panel PCR results obtained in human primary hepatocytes transduced with Nanoblades. Right-panel (top), FACS analysis of CD34+ cells purified from human cord-blood. Bottom, genome editing at the *MYD88* locus assessed by PCR in untreated and Nanoblades-treated CD34+ cells.

Similarly to hiPSCs, mouse bone marrow (BM) cells can be collected and differentiated *in vitro* into various hematopoietic cell types such as macrophages (Bone marrow derived macrophages or BMDMs) and dendritic cells. Efficient genome editing of specific genes in BM cells would therefore allow for the corresponding pre-existing protein to be degraded during differentiation and obtain a functional knockout. To test this hypothesis, BM cells obtained from GFP transgenic mice^7^ were transduced with Nanoblades programmed with a sgRNA targeting the *GFP* coding sequence. 6 hours after transduction, cells were washed and incubated in presence of Macrophage colony-stimulating factor (MCSF) for 1 week. After this, cells were collected to monitor GFP levels by fluorescence microscopy, FACS and genome editing by T7 endonuclease assay (Figure 2b). We consistently obtained close to 75% reduction of GFP expression as measured by FACS analysis and around 60 to 65% genome editing at the *GFP* locus as measured by T7 endonuclease assays (Figure 2b). Importantly, genome editing through Nanoblades did not affect the capacity of BMDMs to respond to LPS as their cytokine expression remains identical to that of untreated control cells (Figure 2b bottom right panel). Nanoblades can therefore be used to inactivate genes in bone marrow cells and study their function in differentiated cells.

Nanoblades efficiency was also tested in a third primary target, primary human hepatocytes. These cells represent a major interest in research and gene therapy due to their proliferative potential and their capacity to colonize and regenerate fully functional tissues. For this, Nanoblades programmed with two sgRNAs targeting the human *MYD88* gene (Figure 2c, left panel) were incubated with primary human hepatocytes for 1 hour. Cells were then grown for 24 hours before extracting their genomic DNA and amplifying the region of *Myd88* flanking the two targeted sites (Figure 2c, right panel). Our results show significant genome editing in these cells and thus indicate that Nanoblades can be used to mediate specific gene deletion in primary human hepatocytes in a transgene deficient manner.

Based on the high efficiency of genome editing in mouse BM cells, we tested whether Nanoblades could also allow genome-editing in human hematopoietic stem cells (HSCs). HSCs are difficult to transduce with VSV-G pseudotyped lentiviral vectors (LVs) because they lack the LDL receptor ^8^. However, LVs pseudotyped with the baboon retroviral envelope glycoprotein (BaEV) have been shown to efficiently transduce HSCs ^9^. We therefore prepared BaEV and VSV G-pseudotyped Nanoblades programmed with two sgRNAs targeting the human *MYD88* gene (Figure 2c) and incubated them with human pre-stimulated CD34+ cells. 48 hours post transduction, cells were collected and genome editing was assessed by PCR using primers flanking the excised sequence. As observed, Nanoblades were also able to induce genome editing in these cells (50% genome editing based on TIDE analysis) thus expanding the catalog of primary cells that can be edited using Nanoblades (Figure 2c).

Taken together, our results indicate that Nanoblades are an efficient delivery system to induce rapid and effective genome editing in murine and human primary cells of high therapeutic value that are notoriously difficult to transfect.

### “All-in-one” Nanoblades for homology directed repair

Precise insertion of genetic material (also known as Knock-in) using CRISPR/Cas9 can be achieved through Homology-Directed Repair (HDR). This occurs when a donor DNA template with sequence homology to the region surrounding the targeted genomic locus is provided to cells together with the Cas9/sgRNA RNP. Based on a previous finding showing that retroviral-particles can be complexed with DNA to allow for virus-dependent DNA transfection ^10^, we decided to test whether Nanoblades could be directly complexed with a DNA template to mediate HDR in target cells. For this, Nanoblades programmed to target a locus close to the AUG start codon of the human *DDX3* gene were complexed to a single-stranded DNA oligomer bearing the FLAG-tag sequence flanked with 46 nucleotide (nt) homology arms corresponding to the region surrounding the start-codon of *DDX3* (Figure 3a, left panel). These “All-in-one” Nanoblades were incubated with HEK-293T cells. 48 hours after transduction, HDR was assessed in the bulk cellular population both by PCR and by Western-blotting (using a FLAG-antibody). As observed (Figure 3a, right panel), cells transduced with “All-in-one” Nanoblades showed incorporation of the FLAG-tag at the *DDX3* locus both genetically and at the level of protein expression. Interestingly, the efficiency of HDR in the bulk population was correlated to the amount of donor DNA template (ranging from 0.01 to 10 pmoles) incubated with Nanoblades (Figure 3a, right panel).

**Figure 3.**
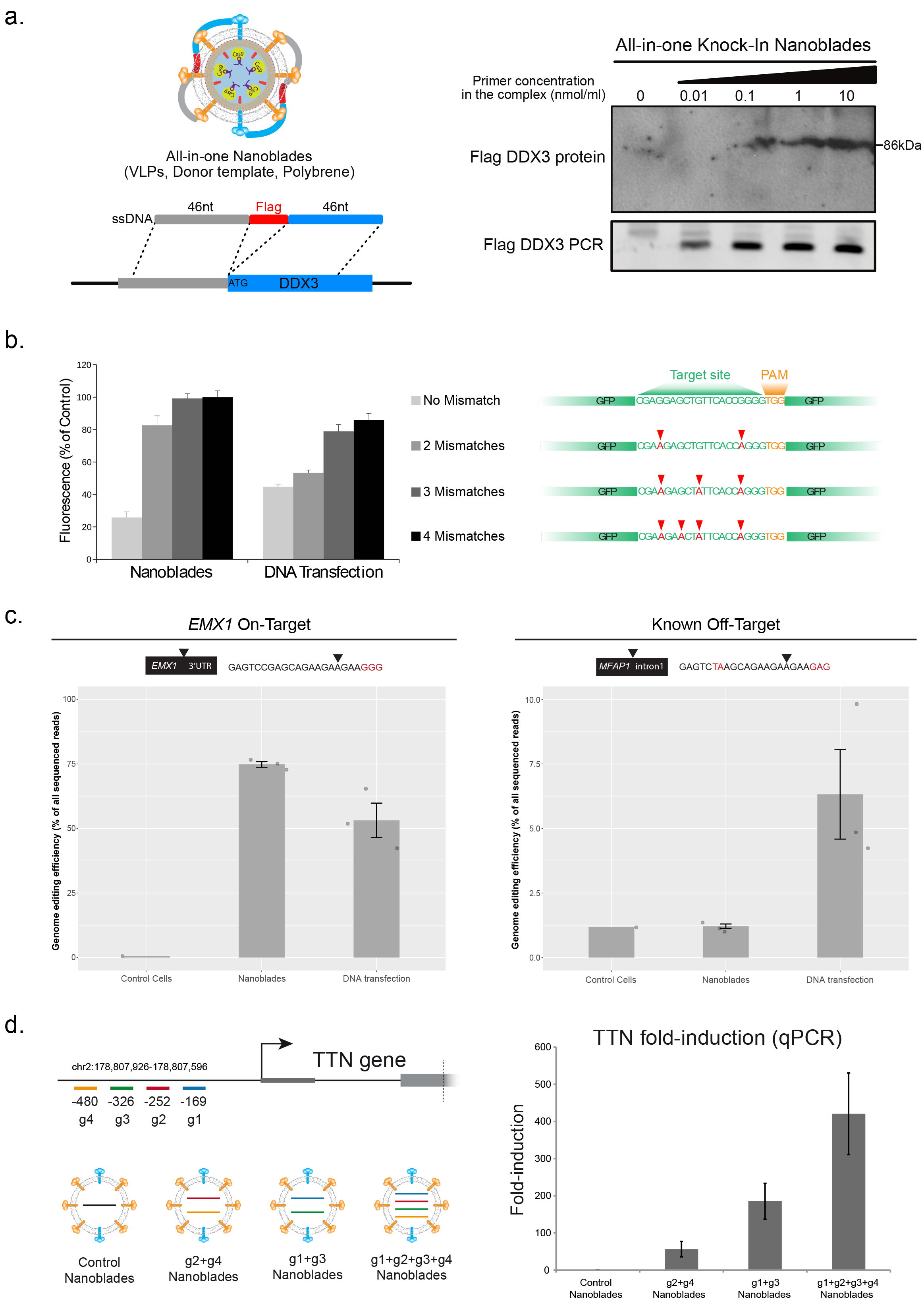
“All-in-one” Nanoblades for knock-in experiments and assessment of Nanoblades off-target activity. **a**. Left panel, Nanoblades targeting human *DDX3* close to its start codon were complexed with a donor ssDNA bearing homology arms to the targeted locus and a Flag-tag sequence in presence of polybrene. Right panel, insertion of the Flag-tag in HEK-293T cells transduced with “all-in-one” Nanoblades complexed with increasing amounts of donor ssDNA assessed by Western-blot using anti-flag antibodies and by PCR. **b**. Left panel, off-target monitoring in immortalized mouse macrophages stably expressing *GFP* transgenes bearing silent mutations in the region targeted by the sgRNA. Right panel, cells were transfected with plasmids coding for Cas9 and the sgRNA or transduced with Nanoblades. GFP expression was measured by FACS 72 hours after transfection/transduction. **c**. Left and right panels, gene-editing at the *EMX1* on-target site and the *MFAP1* intronic off-target site measured by high-throughput sequencing in untreated cells (Control Cells) and cells transduced with *EMX1* Nanoblades (Nanoblades) or transfected with plasmids coding for Cas9 and the EMX1 sgRNA (DNA transfection). **d**. Left panel, position of sgRNAs targeting the promoter of *TTN* and VLPs with different combination of sgRNAs produced for the experiment. Right-panel, TTN mRNA expression levels (normalized to Control) as measured by qPCR in MCF7 transduced with VLPs.

Knock-in assisted by Nanoblades was also obtained inside the AAVS1 locus which has been descrived as a safe harbor for transgene insertion ^11^. For this we designed a dsDNA template of 4 Kb bearing the puromycin resistance gene with homology arms to the AAVS1 locus. After transduction of HEK-293T cells with Nanoblades complexed with this template, single-cell derived clones were selected with puromycin. A PCR-assay revealed that 5 out of 6 puromycin-resistant clones had the puromycin cassette inserted at the AAVS1 locus (Supplementary Figure 3).

Taken together, our results show that Nanoblades can be used for the precise insertion of genetic material through HDR with no requirement of any transfection reagent to introduce the donor DNA template.

### Nanoblades confer low off-target genome-editing

A major concern regarding the use of CRISPR/Cas9-mediated gene editing for gene therapy purposes resides in the potential off-target effects that can occur at genomic loci that are similar in sequence to the original target. Interestingly, several reports have shown that transient delivery of the Cas9/sgRNA complex by injection or RNP transfection generally leads to reduced off-target effects as compared to constitutive expression of Cas9 and sgRNA from DNA transfection experiments ^12^. Since Nanoblades deliver the Cas9/sgRNA complex in a dose-dependent and transient fashion, we tested whether they could also lead to reduced off-target effects when compared to classical DNA transfection. For this, we developed an approach similar to that described by Fu and colleagues ^13^ by creating a series of HEK-293T reporter cell lines transduced with different versions of a *GFP* transgene bearing silent point mutations located in the sgRNA target site (Figure 3b, right panel). These cells were either transfected with plasmids coding for Cas9 and the sgRNA targeting the *GFP* or transduced with Nanoblades programmed with the same sgRNA. 96 hours after transfection/transduction, cells were collected and GFP expression was monitored by FACS (Figure 3b, left panel). As expected, GFP expression from cells bearing the wild-type *GFP* sequence (No Mismatch) was efficiently repressed both after Nanoblades transduction (close to 80% repression) and DNA transfection (close to 60% repression) (Figure 3b, left panel “No Mismatch”). Strikingly, when 2 mismatches were introduced in the target site, Nanoblades were no longer able to efficiently repress GFP expression (20% compared to control) while GFP expression from transfected cells was still reduced to levels similar to that of the *GFP* bearing a perfect match with the sgRNA. Interestingly, the presence of 3 or 4 mismatches totally abolished *GFP* inactivation in Nanoblades-treated cells while cells transfected with the Cas9 and sgRNA plasmids still displayed a mild inhibition of GFP expression (Figure 3b see 3 and 4 Mismatches).

To complement these results, we further tested for genomic off-target effects using the well-characterized sgRNA targeting human *EMX1*. Off-targets for this sgRNA have been extensively studied using T7 endonuclease assays and high-throughput sequencing approaches^14^. We PCR-amplified the *EMX1* locus and one of the previously described *EMX1* genomic off-target loci occurring at the intron of *MFAP1*^14^ in cells treated for 72 hours with Nanoblades programmed with the *EMX1* sgRNA or transfected with a DNA construct coding for Cas9 and the *EMX1* sgRNA. We then assessed genome-editing on each sample by high-throughput sequencing (Figure 3c) ^15^. Editing at the on-target site was efficient in Nanoblade-treated cells (75% in average) and to a less extend in cells transfected with the DNA coding for Cas9 and the sgRNA (53% in average) (Figure 3c, left panel). As expected, small INDELs (insertions and deletions) occurred close to the expected Cas9 cleavage site located 3nt upstream the PAM sequence both in Nanoblades-treated and in DNA transfected cells (Supplementary Figure 4). Surprisingly, in spite of the higher editing efficiency at the on-target site, we could not detect any significant editing at the *MFAP1* off-target site in Nanoblades-treated cells (Figure 3c, right panel). In contrast, cells transfected with the DNA coding for Cas9 and the sgRNA displayed close to 6% editing at the off-target site (figure 3c, right panel) and had INDELs at the expected cut site (Supplementary figure 4).

Taken together, our results indicate that similarly to other protocols that lead to transient delivery of the Cas9/sgRNA RNP, Nanoblades display low off-target effects.

### Targeted transcriptional activation through Nanoblades

Having shown efficient genome editing using Nanoblades loaded with the catalytically active Cas9, we decided to test whether Nanoblades could also deliver Cas9 variants for applications such as targeted transcriptional activation. To this aim, we fused the Cas9-derived transcriptional activator (SP-dCas9-VPR) ^16^ to Gag from MLV and expressed the fusion protein in producer cells together with a control sgRNA or different combinations of sgRNAs targeting the promoter region of human Titin *(TTN)* as previously described ^16^ (Figure 3d, left panel). Nanoblades loaded with SP-dCas9-VPR were then incubated with MCF-7 cells and induction of TTN measured by quantitative PCR (normalized to GAPDH and 18S rRNA expression). As observed (Figure 3d, right panel), when 2 different sgRNAs were used in combination, TTN transcription was stimulated from 50 to 200 fold compared to the control situation. Interestingly, when combining the 4 different sgRNAs in a single VLP, we obtained up to 400 fold transcription stimulation of TTN after 4 hours of transduction. Similar results were obtained at longer time points (Data not shown). Our results therefore suggest that in spite of the large molecular size of the SP-dCas9-VPR (predicted at 224kDa alone and 286kDa when fused to MLV Gag), neither its encapsidation within VLPs nor its delivery and function within target cells are impaired.

### Nanoblade-mediated transduction of zygotes for generating transgenic mice

CRISPR/Cas9 has been extensively used to generate transgenic animals through microinjection of zygotes with DNA coding for Cas9 and the sgRNA or with the synthetic sgRNA and a Cas9 coding mRNA or directly with the preassembled Cas9/sgRNA RNP ^17^. However, all these options require injection into the pronucleus or the cytoplasm of zygotes, which can significantly impact their viability. Moreover, in some species, pronucleus and even cytoplasmic microinjection can be technically challenging.

Because Nanoblades are programmed to fuse their membranes with cellular ones similarly to retroviral and lentiviral vectors, we reasoned that they could also transduce murine zygotes without requiring intracellular microinjection. To test this hypothesis, VLPs loaded with the mCherry protein (instead of Cas9) were produced and injected in the perivitelline space of 1-cell embryos (Figure 4a, top panel). The injected embryos were harvested after 24 hours (2-cell embryos) or 80 hours (blastocysts) for fluorescence analysis (Figure 4a, bottom panel) indicating that VLPs remain mainly in the perivitelline at early stages and subsequently deliver their content into most cells of the developing embryo. Based on these results, Nanoblades programmed with a sgRNA targeting the YFP coding sequence were produced and injected in the perivitelline space of single-cell embryos obtained from YFP transgenic mice ^18^ Transduced embryos were implanted into pseudopregnant females and carried to term. Importantly, we did not observe any significant effect of perivitelline injection on embryo viability. Screening of YFP editing performed by T7 endonuclease assay revealed that 4 out of 8 founder animals displayed significant YFP editing (Figure 4b, top panel) that ranged between 5 and 25% based on Tide analysis of the PCR amplicons and high-throughput sequencing (Data not shown). When 3 out of the 4 founder animals were mated with wild-type mice, we found that the mutant edited allele could be successfully transmitted to the next generation as shown by T7 endonuclease assays on the YFP gene (Figure 4b, bottom panel). Surprisingly, even though we were expecting each F1 individual to bear a single type of mutation in the YFP locus (since only one allele should be transmitted from the founder animal), Tide analysis of the YFP locus from F1 mice indicated that each individual beared a large diversity of YFP mutations (Data not shown). These results suggests that the initial mice strain contains multiples copies of the YFP transgene (a typical feature in transgenic mice obtained from DNA injection procedures). This was further confirmed by quantitative PCR experiments performed on genomic DNA obtained from the parental mouse strain, which indicated an average of 35 copies of the YFP per cell (Data not shown).

To further confirm the ability of Nanoblades to mediate genome editing in mouse embryos, we designed a sgRNA targeting the loxP sequence that could mimic the action of the Cre recombinase by removing a loxP flanked cassette (Figure 4c, left panel). As a first test, these Nanoblades were tested in primary bone marrow cells derived from R26R-EYFP transgenic mice bearing a single-copy of the YFP transgene under control of a “lox-stop-lox” cassette ^19^. As a control, we prepared VLPs loaded with Cre (instead of the Cas9 protein) that should efficiently remove the loxP cassette to induce YFP expression. As expected, Cre-loaded VLPs were able to induce YFP expression in 90% of the bone marrow cells derived from the R26R-EYFP mice (Figure 4c, top right panel). Similarly, Nanoblades programmed with the loxP targeting sgRNA were also able to excise the “lox-stop-lox” cassette and induce YFP expression in 26% of the cells thus validating the sgRNA design (Figure 4c, top right panel). Anti-loxP sgRNA loaded Nanoblades were then injected in the perivitelline space of heterozygous R26R-EYFP 1-cell embryos which were then implanted into pseudopregnant females and carried to term. In this case, 1 out of 14 founder animals was YFP positive under ultraviolet (UV) light and displayed efficient excision of the “lox-stop-lox” cassette as confirmed by PCR (Figure 4c, bottom left panel). Consistent with our previous results, the F1 progeny obtained after mating the loxed F0 mouse with a wild-type mouse contained the “loxed” version of the YFP allele and displayed YFP expression in tails and muscle fibers (Figure 4c, bottom right panel), indicating efficient transmission of the loxed allele from the F0 founder to its progeny.

**Figure 4.**
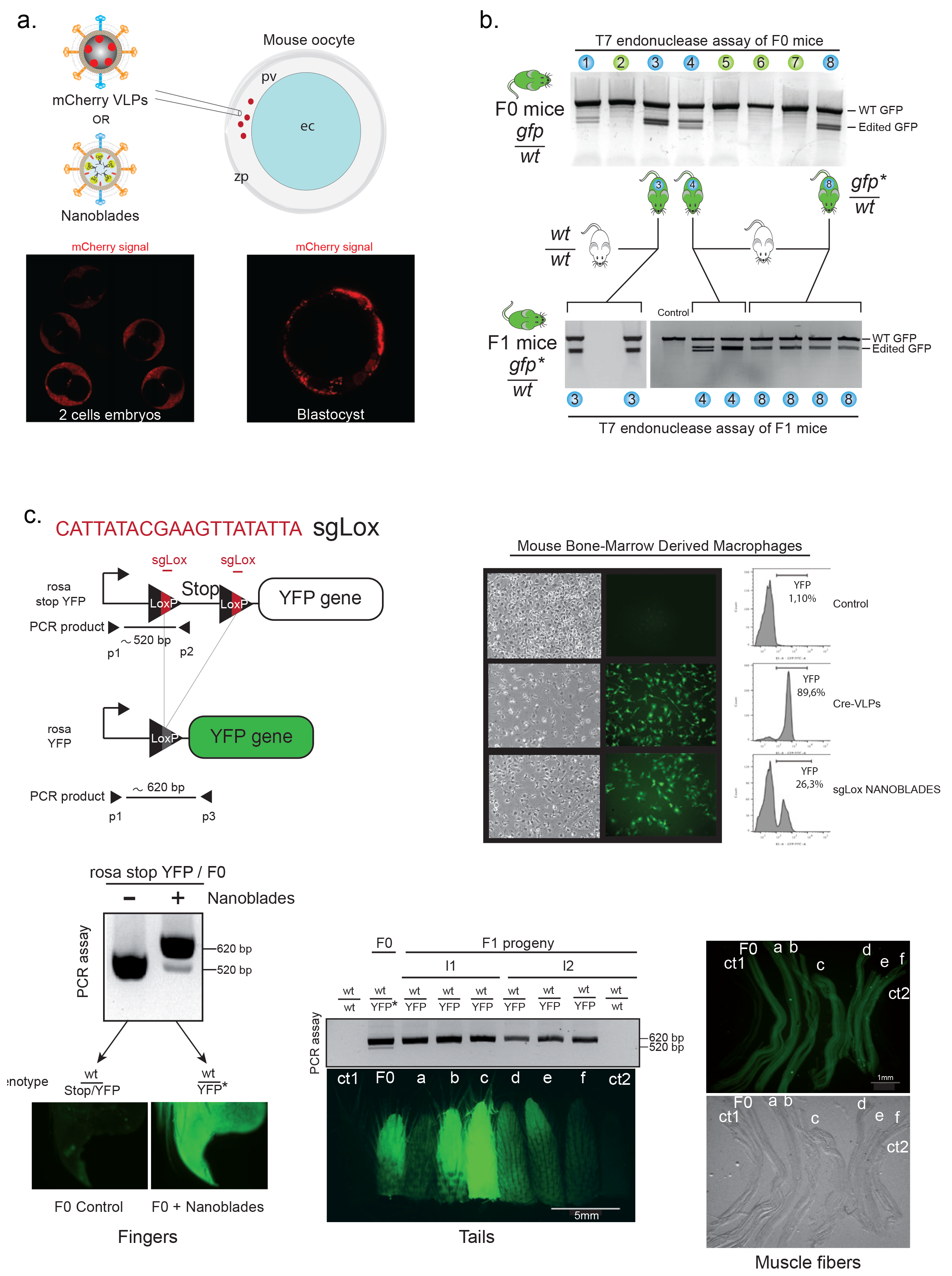
Generation of transgenic mice using Nanoblades. **a**. Top panel, scheme describing injection of mCherry VLPs or Nanoblades in the perivitelline space of mouse 1cell embryos. Bottom panel, fluorescence microscopy of mouse oocytes injected with mCherry VLPs at the single-cell stage. **b**. Top panel, T7 endonuclease assay of F0 GFP-mice obtained after perivitelline injection of zygotes with Nanoblades targeting the *GFP* coding sequence. Bottom panel, T7 endonuclease assay of F1 heterozygous mice obtained by crossing positive T7 endonuclease signal F0 mice (Mice number 3, 4 and 8) with wild-type mice. **c**. Top left panel, scheme describing the inducible YFP lox cassette in the ROSA locus of BL57/CJ6. Top right panel, Fluorescence microscopy and FACS analysis of BMDMs derived from mouse bone marrow cells transduced with Nanoblades programmed with anti-LoxP sgRNAs. Bottom left panel, genetic (PCR) and phenotypic (fluorescence microscopy) analysis of an F0 mouse obtained after injection of ROSA stop YFP mouse oocytes with Nanoblades programmed with anti-LoxP sgRNAs. Bottom middle and right panel, Genetic (PCR) and phenotypic (fluorescence miscroscopy) analysis of the tail and muscle fibers from F1 mice derived from the Nanoblade-treated F0 rosa stop YFP individual.

Taken together, Nanoblades can represent a viable alternative to classical microinjection experiments for the generation of transgenic animals, in particular for species with fragile embryos or with poorly visible pronuclei.

## Discussion

Gene alteration by CRISPRs or other gene editing systems should ideally be achieved in cells or organism by a fast and precise method to limit toxicity of the *modus operandi* and possible off-target effects due to a sustained expression of effectors. In this regard, extensive efforts have been recently described to vehicle Cas9/sgRNA RNPs in cultured cells and *in vivo* by non-coding material including Nanocarriers^20^, optimized transfection reagents ^12^, or lentivirus-derived particles ^21^.

This work describes and characterizes original Virus-Like particles to efficiently vectorize the CRISPR/Cas9 system into primary cells embryos, animal and embryos. These non-coding agents-we called herein Nanoblades-incorporate the Cas9 endonuclease into their internal proteic structure. Beyond delivery of Cas9/sgRNA complexes, we show that Nanoblades can be complexed with reparation templates to mediate homologous-recombination-based knock-in in cultured cells. The molecular basis of this technology is the fusion of Cas9 from *Streptococcus pyogenes* to Gag from Murine Leukemia Virus. Expressed with other components of viral assembly and construct encoding gRNA(s), this molecule can bind sgRNAs into producer cells, forms RNP complexes and cohabits with Gag and Gag-Pol within particles. We indeed show that robust packaging of sgRNAs into Nanoblades depends on their interaction with Gag::Cas9 (Supplementary Figure 1d). RNA-seq analysis of Nanoblades targeting the *GFP* gene revealed that gRNA-GFP was one of the most represented species detected in particles after small cellular RNAs, illustrating that packaging of sgRNAs through Cas9 is an active and efficient process (Supplementary Figure 2).

Similarly to cell-derived particles including most viral vectors, Nanoblades incorporate diverse biomaterials from producer cells, including proteins and cellular RNAs that could be virtually responsible for the transmission of undesired effects. To characterize Nanoblades composition, Mass Spectrometry was performed and give an overview of main proteins transmitted in recipient cells (Supplementary Figure 2). Among the gene ontology terms pointed out by this analysis, plasma membrane terms were particularly enriched which is consistent with the vesicular nature of Nanoblades. As previously described for retroviral-VLPs^22^, characterization of RNA content revealed that Nanoblades contain thousands of individual cellular mRNA species, most of these being encapsidated stochastically, in proportion to their abundance in the producer cell. We found that transcripts overexpressed for production purposes (GAG, VSV-G‥) represent less than 0,4% ot Nanoblades RNAs (Supplementary Figure 2) supporting the notion that their expression in recipient cells is null or marginal. While we cannot exclude the fact that VLPs may be responsible for some cellular responses-depending on recipient cell types-, efficient doses of Nanoblades were globally harmless for most primary cells we tested. In our effort to exploit the retroviral nature of Nanoblades, we explored diverse pseudotyping options (Supplementary Figure 5) and finally focussed on the use of an original mixture of two envelopes (VSV-G plus BRL), a recipe that we have optimized (Supplementary Figure 5) and which systematically displayed the best cleavage results in most recipient cells. Since efficient Nanoblades can be produced with envelopes fusing at the plasma membrane or in endosomes, we suppose that both entry mechanisms can support the delivery of active RNPs and that both of them are used by VSV-G/BRL Nanoblades. Depending on the cellular target, it may be imaginable to equip surface of VLPs with envelopes from Measles virus ^23^, influenza virus ^24^ or other targeting systems ^25, 26^ to restrict or improve Cas9 delivery to certain cell types.

Next generation-Nanoblades may also benefit from the continual evolutions of Cas9-derivatives that can support fusion with Gag from MLV (Figure 3) and could be adapted to other gene-editing targetable nucleases like Cpf1 nucleases ^27^. We also noted that Nanoblades can be engineered to accommodate other proteins/RNAs in addition to Cas9-RNPs and serve as multifunctional agents. Nanoblades capable of delivering both Cas9-RNPs and a reverse-transcribed template that can serve for reparation by homologous-recombination could be envisioned. As illustrated in our work, Nanoblades production can be easily customized by replacement or association of constructs which encode sgRNAs into producer cells. Up to 4 different sgRNAs can be multiplexed in dCas9-VPR Nanoblades (Figure 3) and we commonly used couples of sgRNAs within a single preparation to mediate deletions. We noted that this *modus operandi* is preferable to the pooling of preparations of two particle types, each of them programmed with a single sgRNA. Multiplexing of sgRNAs may also allow the introduction of an additional sgRNA targeting a specific gene that will allow selection of cells efficiently edited by Nanoblade-mediated CRISPR ^28^.

This versatility allows any laboratory equipped with BSL2 facilities to generate its own batches of particles. Beyond cell lines, our VLP-based technique provides a powerful tool to mediate gene editing in primary cells including macrophages, hiPSCs, human hematopoietic progenitors and primary hepatocytes. We have shown that Nanoblade injection into the perivitelline space of mouse-zygotes was particularly harmless for the recipient cells since none of the injected zygotes were affected in their development after treatment. Generation of transgenic animals upon perivitelline space injection of VLPs could be adapted to other species, including larger animals for which the number of zygotes is limitant.

Considering the examples provided in our work, we believe that Nanoblade technology will facilitate gene editing for therapeutical purposes and the rapid generation of primary cell-types harboring genetic diseases, humanized-liver mouse models and transgenic animal models.

## Methods

### Cell culture

Gesicle Producer 293T (Clontech 632617), U2OS cells and primary human fibroblasts (Coriell Institute, GM00312) were grown in DMEM supplemented with 10% fetal calf serum (FCS).

Human induced Pluripotent Stem Cells (hiPSCs) were obtained and cultured as described in ^29^.

Bone marrow-derived Macrophages (BMDMs) were differentiated from bone marrow cells obtained from wild-type C57BL/6 mice. Cells were grown in DMEM supplemented with 10% FCS and 20% L929 supernatant containing Macrophage colony-stimulating factor (MCSF) as described in ^30^. Macrophages were stimulated for the indicated times with LPS (Invivogen) at a final concentration of 100 ng/ml.

### CD34+-cell sample collection, isolation and treatment with Nanoblades

Cord blood (CB) samples were collected in sterile tubes containing the anti-coagulant, citrate-dextrose (ACD, Sigma, France) after informed consent and approval was obtained by the institutional review board (Centre international d’infectiologie (CIRI), Lyon, France) according to the Helsinki declaration. Low-density cells were separated over, Ficoll-Hypaque. CD34+ isolation was performed by means of positive selection using magnetic cell separation (Miltenyi MACs) columns according to the manufacturer’s instructions (Miltenyi Biotec, Bergisch Gladbach, Germany). Purity of the selected CD34+ fraction was assessed by FACS analysis with a phycoerythrin (PE)-conjugated anti-CD34 antibody (Miltenyi Biotec, Bergisch Gladbach, Germany) and exceeded 95% for all experiments. Human CD34+ cells were incubated for 18-24h in 24-well plates in serum-free medium (CellGro, CellGenix, Germany) supplemented with human recombinant: SCF (100ng/ml), TPO (20ng/ml), Flt3-L (100ng/ml) (Myltenyi, France). 5×104 prestimulated CD34+-cells were then incubated with nanoblades in 48-well plates in serum-free medium.

### Plasmids

SP-dCas9-VPR was a gift from George Church (Addgene plasmid # 63798). Lenti CRISPR was a gift from F. Zhang (Addgene plasmid #49535). The GagMLV-CAS9 fusion was constructed by sequential insertions of PCR-amplified fragments in an eukaryotic expression plasmid harboring the human cytomegalovirus early promoter (CMV), the rabbit Beta-globin intron and polyadenylation signals. The MA-CA-NC sequence from Friend Murine Leukemia virus (Accession Number :M93134) was fused to the MA/p12 protease-cleavage site (9 aa) and the Flag-nls-spCas9 amplified from pLenti CRISPR.

### sgRNA design and sequences (+PAM)

sgRNAs targeting *MYD88, DDX3, GFP, Hpd* and the *LoxP* sequence were designed using CRISPRseek ^31^.

Human rDNA: 5’ CCTTCTCTAGCGATCTGAGagg 3’
Human *EMX1:* 5’ GAGTCCGAGCAGAAGAAGAAggg 3’
Human *MYD88* #1: 5’ GAGACCTCAAGGGTAGAGGTggg 3’
Human *MYD88* #2: 5’ GCAGCCATGGCGGGCGGTCCtgg 3’
*GFP:* 5’ CGAGGAGCTGTTCACCGGGGtgg 3’
Human *DDX3:* 5’ AGGGATGAGTCATGTGGCAGtgg 3’
Mouse *Hpd*: 5’ GAGTTTCTATAGGTGGTGCTGGGTGggg 3’
Human *TTN*-169: 5’ CCTTGGTGAAGTCTCCTTTGagg 3’
Human *TTN*-252: 5’ ATGTTAAAATCCGAAAATGCagg 3’
Human *TTN*-326: 5’ GGGCACAGTCCTCAGGTTTGggg 3’
Human *TTN*-480: 5’ ATGAGCTCTCTTCAACGTTAagg 3’
*Human AAVS1:* 5’ ACCCCACAGTGGGGCCACTAggg 3’
*LoxP:* 5’ CATTATACGAAGTTATATTAagg 3’

### Production of Nanoblades

Nanoblades were produced from transfected Gesicles Producer 293T Cells plated at 5×10_6_ cells /10-cm plate 24 hours before transfection with the JetPrime reagent (Polyplus). Plasmids encoding the GagMLV-CAS9 fusion (1.7µg), Gag-POLMLV (2.8µg), gRNA expressing plasmid(s) (4.4µg), VSV-G (0.4µg), the Baboon Endogenous retrovirus Rless glycoprotein (BaEVRless) ^9^ (0.7µg) were cotransfected and supernatants were collected from producer cells after 40 hours. For production of serum-free particles, medium was replaced 24 hours after transfection by 10ml of Optimem (Gibco) supplemented with penicillin-streptomycin. Nanoblade-containing medium was clarified by a short centrifugation (500 xg 5 min) and filtered through a 0,8µm pore-size filter before ultracentrifugation (1h30 at 96,000 xg). Pellet was resuspended by gentle agitation in 100µl of cold 1X PBS. Nanoblades were classically concentrated 100-fold. X-Nanoblades referred as Nanoblades loaded with gRNA(s) targeting the x-gene.

To dose Cas9 packaged into particles, Nanoblades or recombinant Cas9 (New England Biolabs) were diluted in 1X PBS and serial dilutions were spotted onto a Nitrocellulose membrane. After incubation with a blocking buffer (nonfat Milk 5%w/v in TBST), membrane was stained with a Cas9 antibody (7A9-3A3 clone, Cell signaling) and revealed by a secondary antiMouse antibody coupled to horseradish peroxidase. Cas9 spots were quantified by Chemidoc touch imaging system (Biorad).

### Combination of Nanoblades with ssDNA and dsDNA

ssDNA (DDX3): 15µl of concentrated DDX3-Nanoblades (2µM Cas9) were mixed with 10µl of 1X PBS containing 8µg/ml of polybrene supplemented with 5µl of each dilutions of the Flag-DDX3 primer, best results being obtained with the higher concentration (5µl of primer at 100pmol/µl). This ‘all-in-one’ complex was incubated 15 min at 4°C and the 30µl were added onto the medium (400µl+polybrene 4µg/ml) of HEK-293T cultivated in a 12-well-plate (200,000 cells plated the day before). After two hours, the transduction medium was supplemented with 1ml of DMEM 10% FCS. 40 hours after VLP-treatment, cells were passaged for amplification and analysis of the genetic insertion of the flag sequence upstream the DDX3 gene and Western-blot analysis were performed 72 hours later. Sequence of the Flag-DDX3 primer (HPLC-purified): 5’-ACTCGCTTAGCAGCGGAAGACTCCGagTTCTCGGTACTCTTCAGGGATGGACTACAAGGACGACGATGACAAGagTCATGTGGCAGTGGAAAATGCGCTCGGGCTGGACCAGCAGGTGA-3’

dsDNA (AAVS1): 20ul of concentrated nanoblades were complexed with 500ng of dsDNA in a total volume of 60 microliter of PBS with polybrene at a final concentration of 4ug/ml. After 15 minutes of incubation on ice, complexes were used to transduce HEK293T cells plated the day before at 100000 cells in a 24-well plate in medium supplemented with polybrene (4ug/ml). Two days latter cells were trypsined and replated in a 10cm dish before Puromycin selection (0,7ug/ml). Single-cell derived clones were next isolated and cultivated in a 12w plates before PCR analysis performed on genomic DNAs (500 ng). Primers used were:

AAVS1forward:5’-TCCTGAGTCCGGAC CACTTT GAG-3’
Puromycin reverse:5’-GATCCAGATCTGGTGTGGCGCGTGGCGGGGTAG-3’
EMXgene forward 5’-TTCTCTCTGGCCCACTGTGTCCTC-3’
EMXgene reverse 5’-AGCCCATTGCTTGTCCCTCTGTCAATG-3’.

### T7 endonuclease assay

Genomic DNA was extracted from VLP-treated cells using the Nucleospin gDNA extraction kit (Macherey-Nagel). 150ng of genomic DNA was then used for PCR amplification. PCR products were diluted by a factor 2 and complemented with Buffer 2 (New England Biolabs) to a final concentration of 1X. Diluted PCR amplicons were then heat denatured at 95°C and cooled down to 20°C with a 0.1°C/second ramp. Heteroduplexes were incubated for 30 minutes at 37°C in presence of 10 units of T7 Endonuclease I (NEB). Samples were finally run on a 2.5% agarose gel or on a BioAnalyzer chip (Agilent) to assess editing efficiency.

### Immunofluorescence, antibodies and imaging

Cells were fixed in 1X PBS supplemented with 4% of paraformaldehyde (PFA) for 20 min, washed three times with 1X PBS and permeabilized with 0.5% Triton X-100 for 4.5 min. Cells were incubated with primary antibodies overnight at 4°C. Primary antibodies used are: rabbit yH2AX (1:1000; Abcam 81299) and mouse RNA pol I RPA194 (1:500; Santacruz sc48385). Cells were washed three times in 1X PBS, followed by incubation of the secondary antibodies conjugated to Alexa 488 or 594 used at a 1:1000 dilution (Life Technologies) for 1 hour at room temperature. After three 1X PBS washes, nucleus were stained with Hoechst 33342 at 1µg/ml for 5 min. The coverslips were mounted in Citifluor medium (AF1, Citifluor, London, United Kingdom). Cells were observed under a Leica DM6000. At least 100 cells were counted in each indicated experiment. Averages and standard deviation values were obtained from three independent biological replicates.

### Mouse experiments

All experiments were performed in accordance with the European Union guidelines for approval of the protocols by the local ethics committee (Authorization Agreement C2EA 15, “Comité Rhône-Alpes d’Ethique pour l’Expérimentation Animale”, Lyon, France.

### Northern-blot of sgRNAs

2µg of total RNA extracted from Nanoblades or Nanoblade-producing cells were run on a 10% acrylamide, 8M Urea, 0.5X TBE gel for 1 hour at 35watts. RNAs were then transferred onto a Nitrocellulose membrane (Hybond Amersham) by semi-dry transfert for 1 hour at 300mA in 0.5X TBE. The membrane was UV-irradiated for 1 minute using a stratalinker 1800 and then baked at 80°C for 30 min. The membrane was then incubated in 50ml of Church Buffer (125mM Na_2_HPO_4_, 0.085% Phosphoric Acid, 1mM EDTA, 7% SDS, 1%BSA) and washed twice in 10ml of Church buffer. The 5’ P32-labeled (1×10_7_cpm total) and heat-denatured ssDNA probe directed against the constant sequence of the guideRNA (sequence of the probe 5’GCACCGACTCGGTGCCACTTTTTCAAGTTGATAACGGACTAGCCTTATTTTAACTTGCTATTTCTAGCTCTA3’) was diluted in 10ml of Church buffer and incubated with the membrane overnight at 37°C. The membrane was washed four times in 50ml of wash buffer (1X SSC + 0.1% SDS) before proceeding to phosphorimaging.

## Acknowledgments

Sequencing was performed by the IGBMC Microarray and Sequencing platform, a member of the ‘France Génomique’ consortium (ANR-10-INBS-0009). We acknowledge the contribution of SFR Biosciences (UMS3444/CNRS, US8/Inserm, ENS de Lyon, UCBL) facilities: Platim and PBES (Celphedia, AniRA), especially M. Teixera for mouse embryos microinjection. We also thank J.F. Henry, N. Aguilera and J.L. Thoumas from the animal facility (PBES, Plateau de Biologie Expérimentale de la Souris, ENS de Lyon), as well A. Ollivier for their technical help in handling mice. We thank N. Gadot (Plateforme Anatomopathologie Recherche, Centre Léon Bérard, F-69373 Lyon, France) for the IHC analyses.

## Author contributions

P.E.M and E.P.R conceived the study and designed most experiments. P.E.M, E.P.R, V.R, A.M, E.L, F.F, E.V, F.L.C, T.S and F.A designed experiments. P.E.M, E.P.R, E.L, V.R, A.M, F.F, T.S, F.A, J.B, E.V, V.M and E.M performed experiments and analyzed data. P.E.M and E.P.R wrote the paper with contributions from all authors.

---

**Supplementary figure 1.**
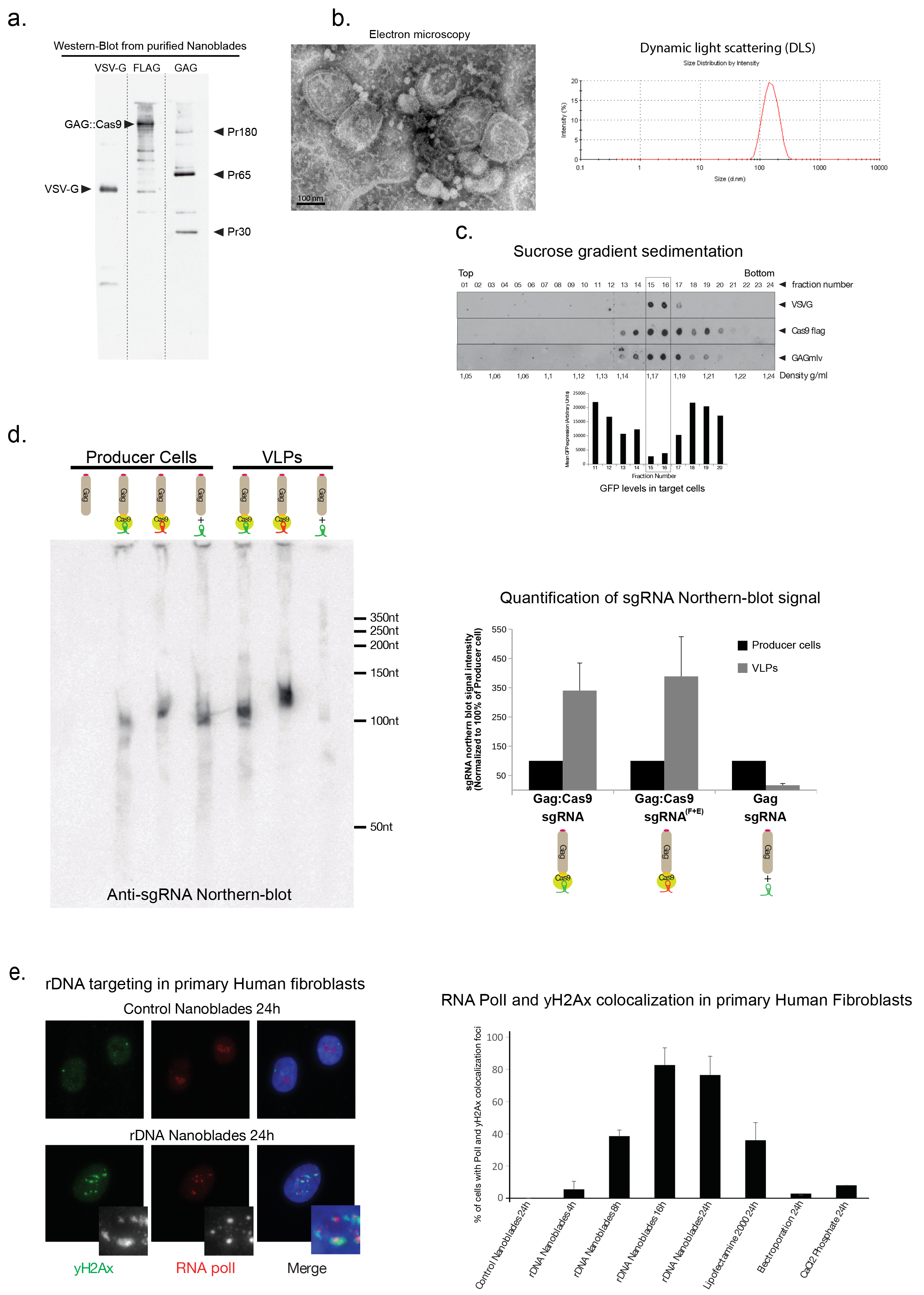
Molecular, structural and biochemical characterization of Nanoblades. **a**. Western blot analysis of proteins from purified Nanoblades using anti VSV-G, anti-Flag and anti-Gag antibodies. **b**. Electron microscopy and dynamic light scattering analysis of purified Nanoblades. **c**. Sucrose sedimentation analysis of Nanoblades targeting the *GFP* coding sequence. Each fraction of the sucrose gradient was analysed by Western-blotting to monitor the presence of VSV-G, Cas9 and Gag. Bottom panel, fractions 11 to 20 were collected and incubated with immortalized mouse macrophages stably expressing GFP. GFP expression was then measured by FACS 96h after transduction. **d**. Left panel, Northern-blot analysis of total RNA extracted from producer cells and purified Nanoblades using a radioactive probe complementary to the conserved region of the sgRNA. From left to right, producers cells expressing Gag only or Gag::Cas9 + sgRNA or Gag::Cas9 + modified sgRNA or Gag + sgRNA and VLPs obtained from cells expressing Gag::Cas9 + sgRNA or Gag::Cas9 + modified sgRNA or Gag + sgRNA. Right-panel, quantification of the Northern-blot signal. **e**. Left panel, immunofluorescence analysis of γ-H2AX (green), RNA pol I (red) in primary human fibroblasts 24 hours after being transduced with control Nanoblades or with Nanoblades targeting ribosomal DNA genes. Right panel, quantification of γ-H2AX and RNA Pol I colocalization foci in primary fibroblasts at different times after Nanoblades transduction or after classical DNA transfection methods.

**Supplementary figure 2.**
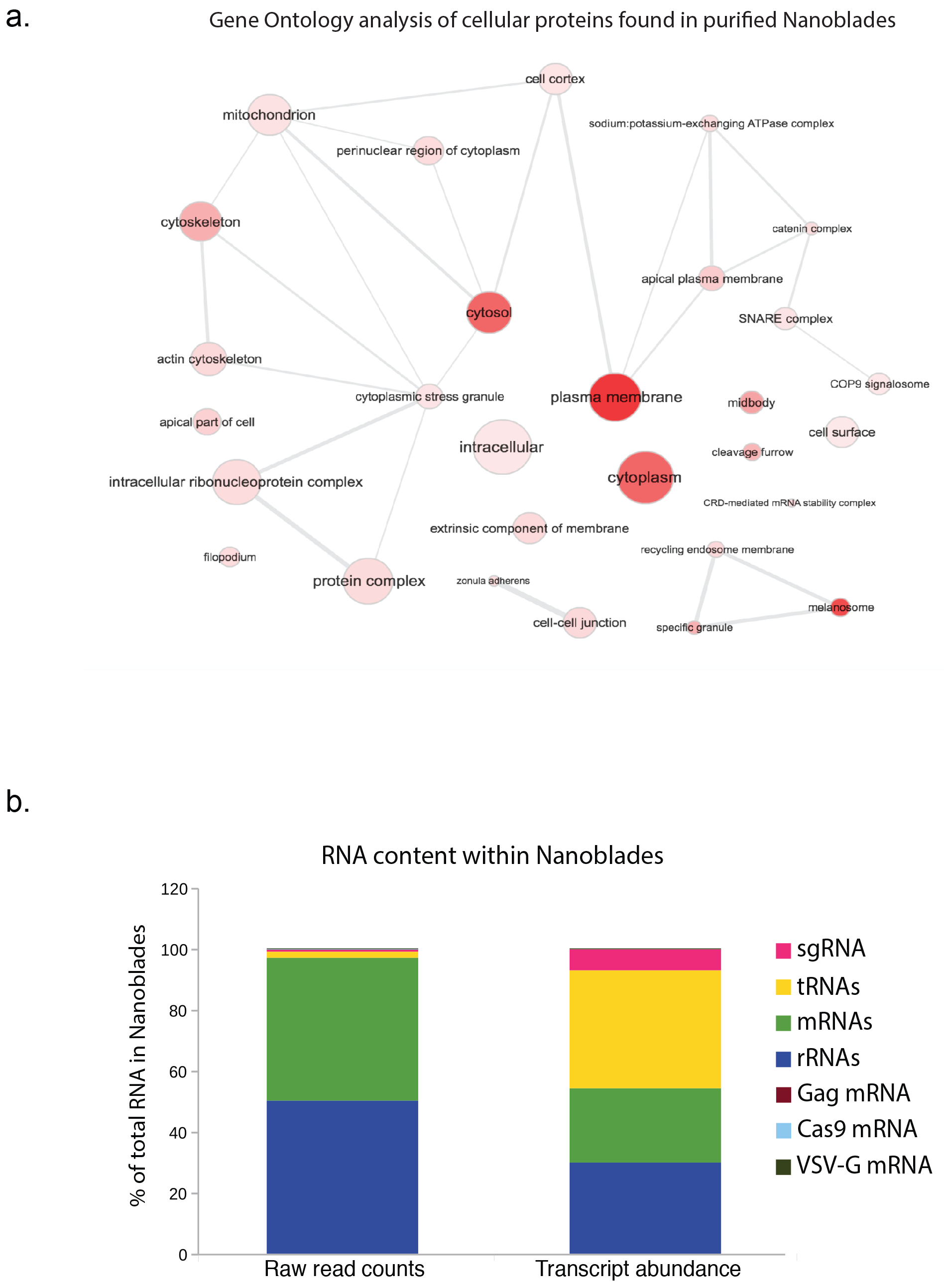
Protein and RNA content of purified Nanoblades. **a**. Gene ontology analysis of proteins identified by Mass Spectrometry in Nanoblades. **b**. Relative quantification of all RNAs found within Nanoblades by high-throughput sequencing.

**Supplementary figure 3.**
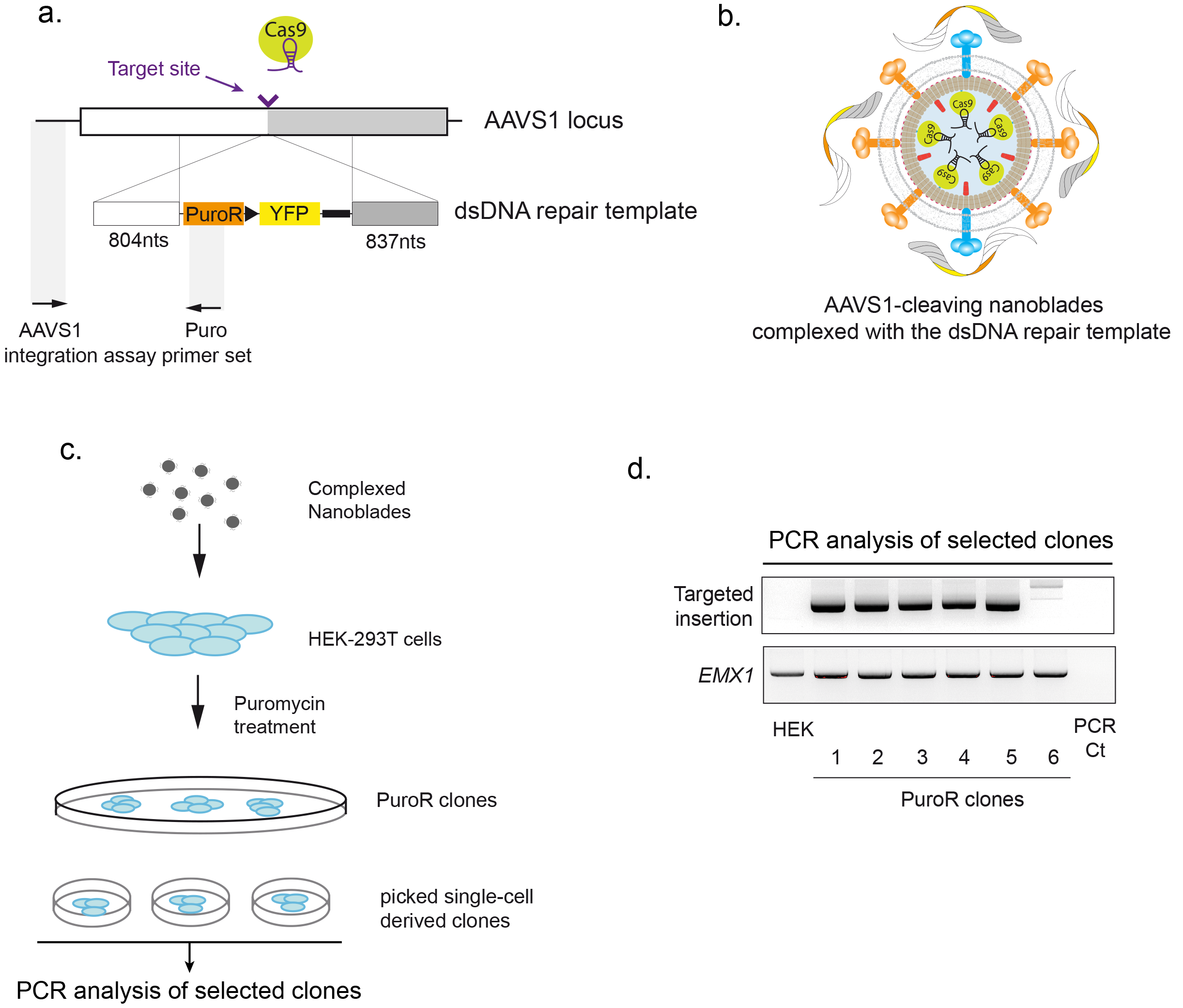
“All-in-one” Knock-in of a puromycin cassette in the *AAVS1* locus. **a**. Scheme of the knock-in strategy and the dsDNA puromycin cassette. **b**. Nanoblades targeting the human *AAVS1* locus complexed with a donor dsDNA puromycin cassette bearing homology arms to the targeted locus in presence of polybrene. **c**. Scheme of the transduction and clonal selection strategy. Briefly, HEK293T cells were transduced with “all-in-one” Nanoblades targeting the *AAVS1* locus and complexed with the dsDNA puromycin resistance cassette. After transduction, cells were incubated with puromycin until individual resistant clones were visible. 6 resistant clones were then isolated to obtain monoclonal cell lines. Targeted insertion of the puromycin cassette in the *AAVS1* locus was then monitored by PCR (using the AAVS1 and Puro oligomers depicted in **a**) upon genomic DNA extraction. **d**. PCR analysis of the *AAVS1* and *EMX1* (control) loci in untreated HEK-293T cells (HEK), in the 6 puromycin resistant clones (PuroR clones) and in water (PCR Ct).

**Supplementary figure 4.**
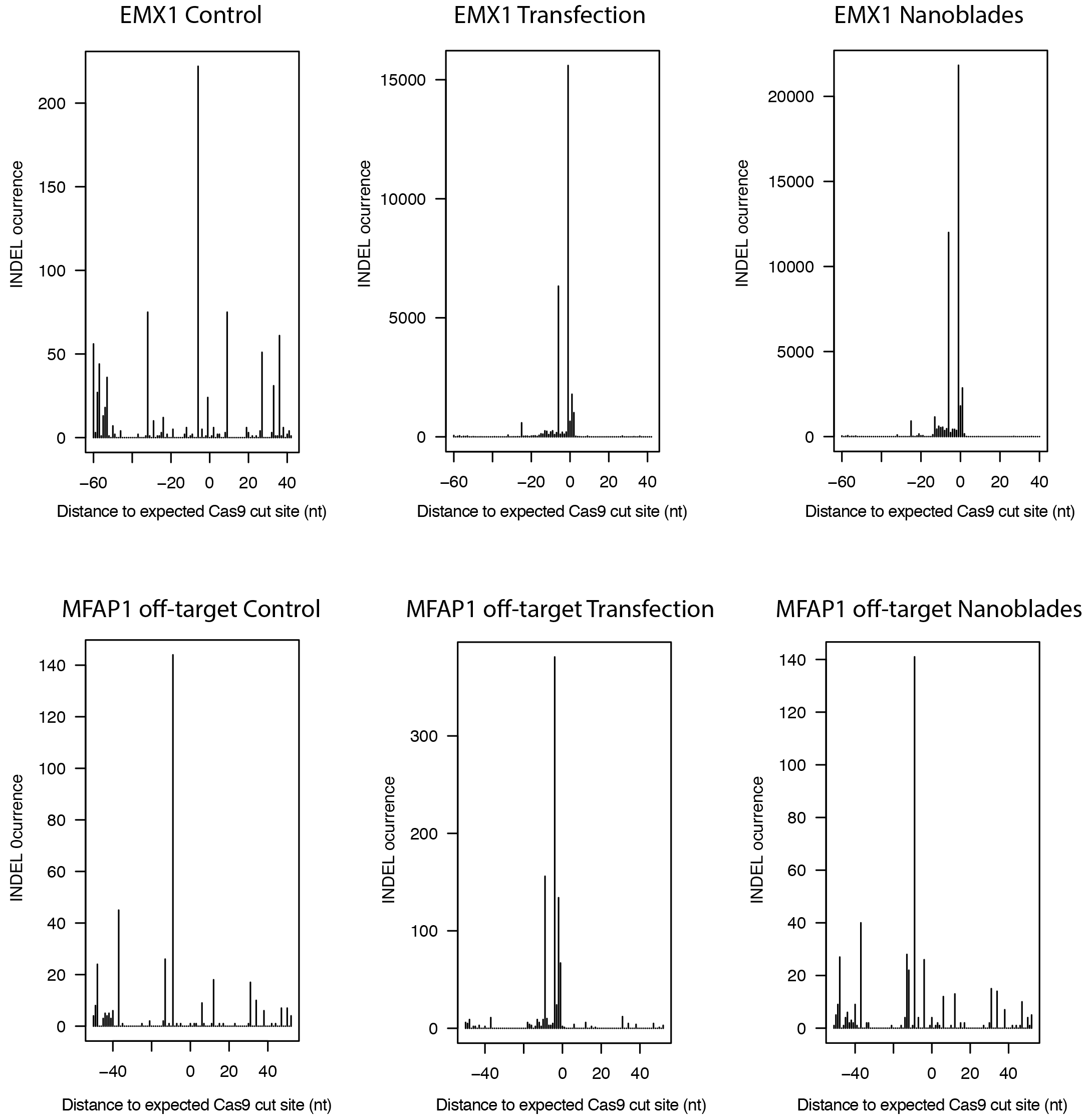
Position of INDELs detected at the EMX1 on-target site and the MFAP1 off-target site. The frequency of INDELs detected in high-throughput sequencing reads were plotted for each position of the EMX1 (Top panels) and MFAP1 (Bottom panels) loci using the expected Cas9 cut site (3nt upstream the PAM site) as the 0 offset position.

**Supplementary figure 5.**
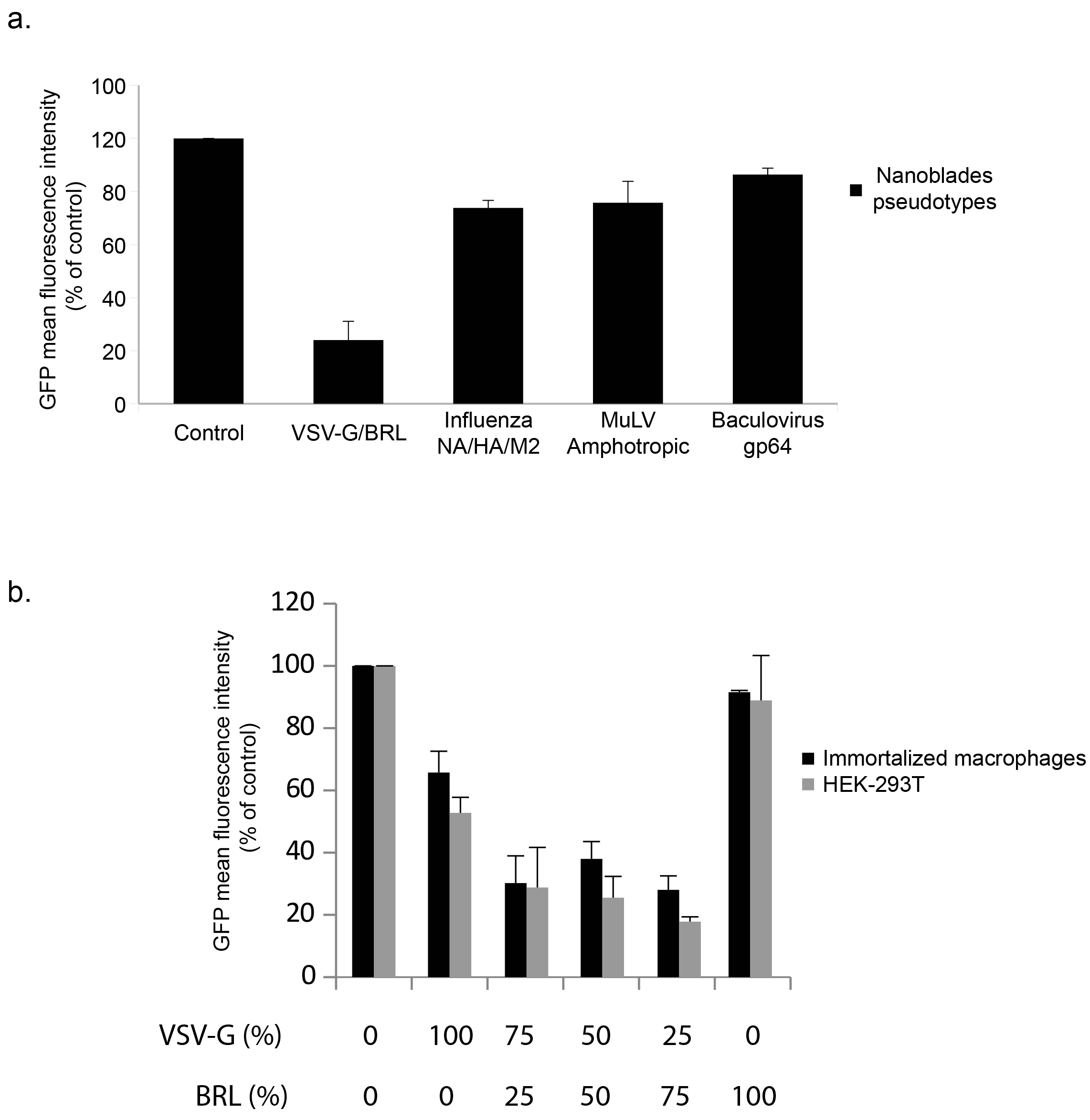
Pseudotyping of Nanoblades with different envelope glycoproteins. **a**. Nanoblades programmed with a GFP targeting sgRNA and pseudotyped with different viral-derived envelope glycoproteins were incubated with immortalized mouse macrophages that constitutively express GFP. 72 hours posttransduction, the mean fluorescence intensity (MFI) was measured by FACS. **b**. Nanoblades programmed with a GFP targeting sgRNA and pseudotyped with different ratios of the VSV-G and BRL envelope glycoproteins were incubated with immortalized mouse macrophages and HEK293T cells that constitutively express GFP. 72 hours posttransduction, the mean fluorescence intensity (MFI) was measured by FACS.

